# A fronto-temporo-parietal network disambiguates potential objects of joint attention

**DOI:** 10.1101/542555

**Authors:** P. M. Kraemer, M. Görner, H. Ramezanpour, P. W. Dicke, P. Thier

**Affiliations:** Department of Cognitive Neurology, Hertie Institute for Clinical Brain Research, University of Tübingen, 72076 Tübingen, Germany.; Graduate School of Neural and Behavioural Sciences, University of Tübingen, 72074 Tübingen, Germany.; International Max Planck Research School for Cognitive and Systems Neuroscience, University of Tübingen, 72074 Tübingen, Germany.; Werner Reichardt Centre for Integrative Neuroscience, University of Tübingen, 72076 Tübingen, Germany.; Center for Decision Neuroscience, Faculty of Psychology, University of Basel, 4055 Basel, Switzerland.

## Abstract

We use the other’s gaze direction to identify her/his object of interest and to shift our attention to the same object, i.e. to establish joint attention. However, gaze direction may not be sufficient to unambiguously identify the object of interest as the other’s gaze may hit more than one object. In this case, the observer must use *a priori* information to disambiguate the object choice. Using fMRI, we suggest that the disambiguation is based on a 3-component network. A first component, the well-known ‘gaze following patch’ in the posterior STS is activated by gaze following per se. BOLD activity here is determined exclusively by the usage of gaze direction and is independent of the need to disambiguate the relevant object. On the other hand, BOLD activity revealing *a priori* information for the disambiguation and starting early enough to this end is confined to a patch of cortex at the inferior frontal junction. Finally, BOLD activity reflecting the convergence of both, a priori information and gaze direction, needed to shift attention to a particular object location is confined to the posterior parietal cortex.

## Introduction

We follow the gaze of others to objects of her/his attention and to shift our attention to the same object, thereby establishing joint attention. By associating our object-related intentions, expectations and desires with the other one, joint attention allows us to develop a *Theory of* (the other’s) *Mind* (TOM). Disposing of a viable TOM is a major basis of successful social interactions ^1,2^ and arguably its absence is at the core of devastating neuropsychiatric diseases such as autism. Human gaze following is geometric^3,4^. This means that we use the other’s gaze vector to identify the exact location of the object of interest. The features of the human eye such as the high contrast between the white sclera and dark iris allow us to determine the other’s eye direction at high resolution^5,6^. However, knowledge of direction is not sufficient to pinpoint an object in 3D. In principle, differences between the directions of the two eyes, i.e. knowledge of the vergence angle, could be exploited to this end. Yet, this will work only for objects close to the beholder as the angle will become imperceptibly small if the objects are outside the confines of peripersonal space. On the other hand, gaze following remains precise also for objects quite far from the other although the gaze vector will in many cases hit more than one object^4^. Hence, how can these objects be disambiguated? We hypothesized that singling out the relevant object is a consequence of recourse to prior information on the objects and their potential value for the other. For instance, let us assume that the day is hot and that the other’s appearance may suggest thirst and the desire to take a sip of something cool. If her/his gaze hit a cool beverage within a set of other objects of little relevance for a thirsty person, the observer might safely infer that the beverage is the object of desire. In this example, gaze following is dependent on prior assumptions about the value of objects for the other. Of course, also the value the object may have for the observer matters. For instance, Liuzza et al. showed that an observer’s appetence to follow the other’s gaze to portraits of political leaders is modulated by the degree of political closeness^7^. If the politician attended by the other was a political opponent of the observer, the willingness to follow gaze was significantly reduced. Also knowing that gaze following may be inadequate in a given situation and that the other may become aware of an inadequate behavior will suppress it^8,9^. However, only assumptions about the object value for the other will help to disambiguate the scene.

Following the gaze of others to a particular object is accompanied by a selective BOLD signal in an island of cortex in the posterior superior temporal sulcus (pSTS), the “gaze-following patch (GFP)”^10–12^. In these studies, the target object could be identified unambiguously by gaze direction as for a given gaze direction the vector hit one object only. Hence, it remained unclear if the GFP helps to integrate the information needed to disambiguate the object choice in case the gaze vector hits more than one object. In order to address this question, we carried out an fMRI study in which the selection of the object of joint attention required that the observer recoursed on another source of information aside from the gaze cue.

## Results

### Behavioral Performance

Our subjects participated in two fMRI experiments. The first one was a localizer task that allowed us to identify two *a priori* defined regions of interest (ROI), the GFP and parietal area hLIP (human LIP). To identify the GFP in the temporal lobe, we compared the BOLD activity evoked by following the gaze of a human avatar to one out of 4 possible target objects (*gaze following*, *gf*) with the activity evoked by using to avatar’s eye color to overtly shift attention to the target sharing this color (*color mapping*, *cm*). A significant *gf* > *cm* contrast delineated a region in the pSTS that matched the coordinates of the *GFP* as known from previous studies ^11,12^. Area hLIP was localized by a significant *cm* > *bl* (baseline) contrast in the parietal lobe. The identified region matched values given elsewhere as well ^13^. The second experiment was a gaze following task, in which the subjects saw a human avatar gazing along one out of four linearly arranged sets of 3 objects each. The objects were selected from two categories, houses and hands. Hands and houses were distributed such that each category was represented by 1 or 2 exemplars. The observers had to follow the avatar’s gaze to a particular object, identified by the conjunction of the avatar’s gaze direction and a verbal instruction that specified the object category relevant in a given trial (cf. Fig. 1 for an illustration). After an initial baseline period, during which the avatar looked straight ahead, subjects observed the avatar making a saccade to one of the four object sets. At the same time, the verbal instruction was delivered. It could either be unambiguous (“house” vs. “hand”, 1/3 of trials each) or remain uninformative (“none”, 1/3 of trials). Depending on the conjunction of gaze direction and instruction three conditions could be distinguished: The *unambiguous condition (ua;* the instruction was informative and there was only one of the verbally specified objects in the set), the *ambiguous-informative condition* (*inf*; two of the objects were in the set) and the *ambiguous-uninformative condition* (*uninf*; the verbal instruction was uninformative, i.e. three possible targets). Participants were asked to use the available information to decide on a target and to communicate their decision by making a saccade to that target 5 s after the avatar’s saccade with the disappearance of the fixation dot serving as go-signal. As their decision had to consider both gaze direction and the context of the verbal instruction we will refer to this task as the *contextual gaze following task*.

**Fig. 1.**
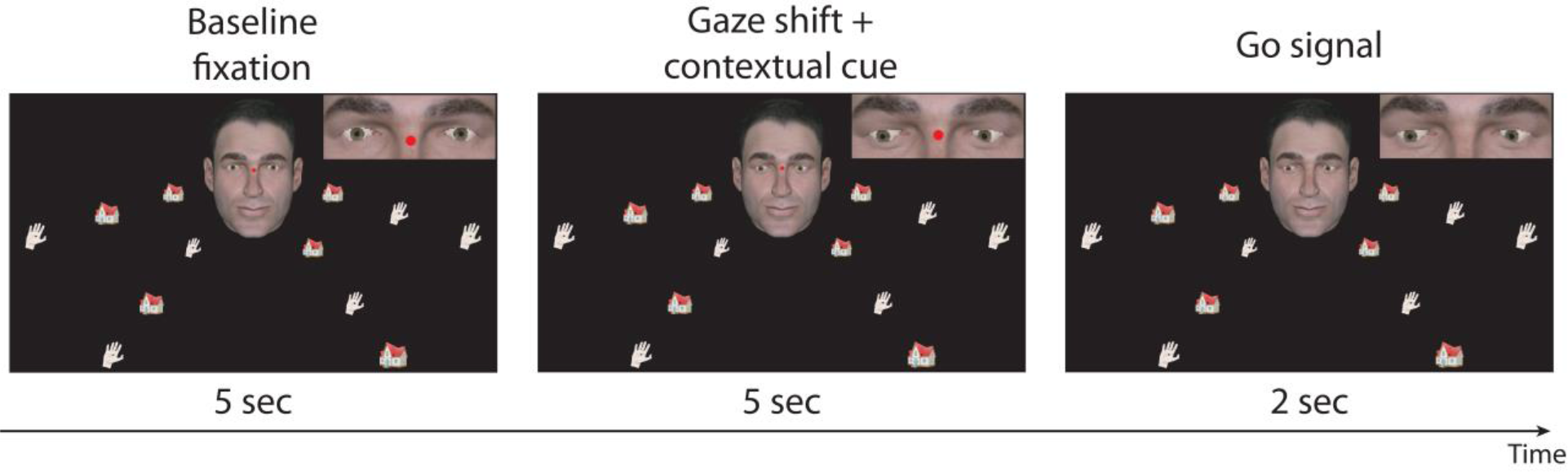
Contextual gaze following task. An avatar appeared in the center of the screen together with four linearly arranged sets of objects (houses and hands). After a baseline fixation period, the portrait’s gaze shifted towards one specific target object simultaneously with an auditory contextual instruction specifying the object class of the target (hand or house) or not, i.e. remaining uninformative (“none”). While maintaining fixation, subjects needed to decide on the target and make a saccade to the chosen target after a go-signal indicated by the disappearance of the fixation dot.

In the localizer task, subjects were able to hit targets reliably and without significant difference between the two conditions (median hit rates: *gf*: 0.94 ± 0.13 s.d.; *cm*: 0.92 ± 0.09 s.d.; *p* = 0.6, two-tailed t-test, *N* = 19, Fig. 2). Using the gaze following performance in the localizer task as reference we estimated the following expected hit rates for the contextual gaze following task: 0.94 for the *unambiguous condition*, 0.94*1/2 for the *ambiguous-informative* and 0.94*1/3 for the *ambiguous-uninformative* condition (Fig. 2). As summarized in Fig. 2, the measured performances matched the estimates in the contextual gaze following task very well (comparison by two-tailed t-tests, n.s.). This result clearly indicates that the probability to identify an object as a target was exclusively determined by the information provided by gaze direction and verbal instruction and not influenced by biases or uncontrolled strategies.

**Fig. 2.**
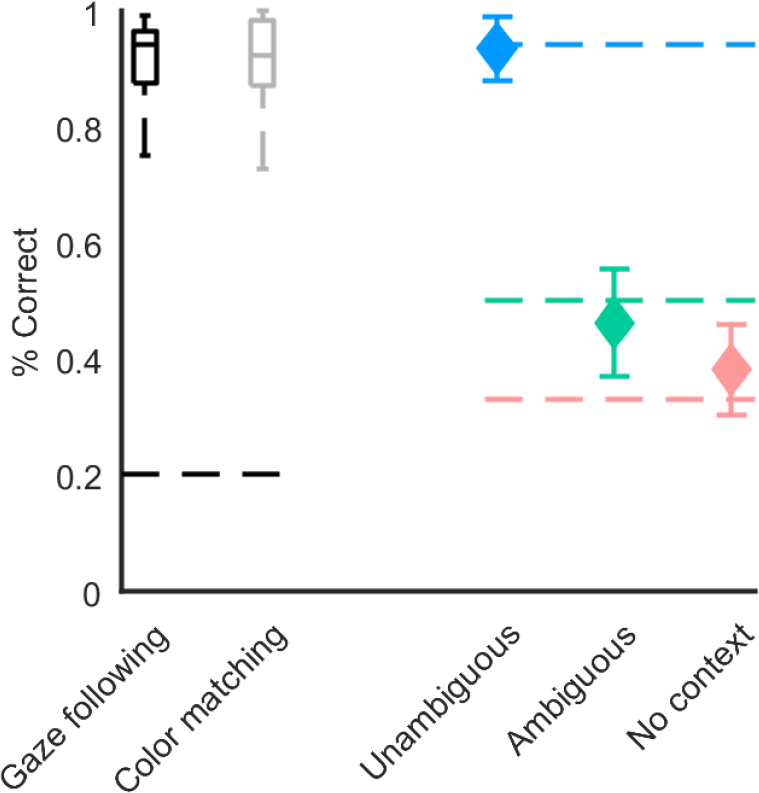
Behavioral performance. Left: Boxplots (black and gray) showing the percentage of correct response in the localizer paradigm (dashed line depicts chance level performance). Right: Plots of correct responses in the contextual gaze following paradigm (weighted mean performance and weighted std, dashed lines depict expected performance).

### Task related brain regions

To localize the GFP we contrasted *gf* with *cm* trials of the first experiment. At the group level (*N* = 19) this contrast yielded a patch of significantly larger activity for *gf* in the pSTS in both hemispheres. The contrast maxima (blue spheres in Fig. 3, upper) were located at *x*, *y*, *z* = −57, −61, −1 in the left and at *x*, *y*, *z* = 48, −67, −1 in the right hemisphere. These locations closely match those known from other studies, visualized as green and cyan spheres for comparison ^11,12^. In addition to the GFP, the *gf* > *cm* contrast was significant in a few more regions, not consistently seen as activated in previous work using the same paradigm (see supplementary material Tab. 1 for a list of all activated regions). Based on the group coordinates of the GFP we tried to localize it in individual subjects by searching for the closest maximum activation which passed a statistical significance threshold (*p* < 0.05, uncorrected) and a cluster size threshold (cluster size >= 6 voxel). Clusters that lay outside of a sphere with a radius of 10 mm centered on the group maximum were excluded (proximity criterion). Under these constraints, we were able to determine individual GFPs for nine subjects in the right and for six subjects in the left hemisphere (white spheres ibid., SD of individual locations: right x, y, z = 5, 5, 3; left x, y, z = 3, 3, 5).

**Fig. 3.**
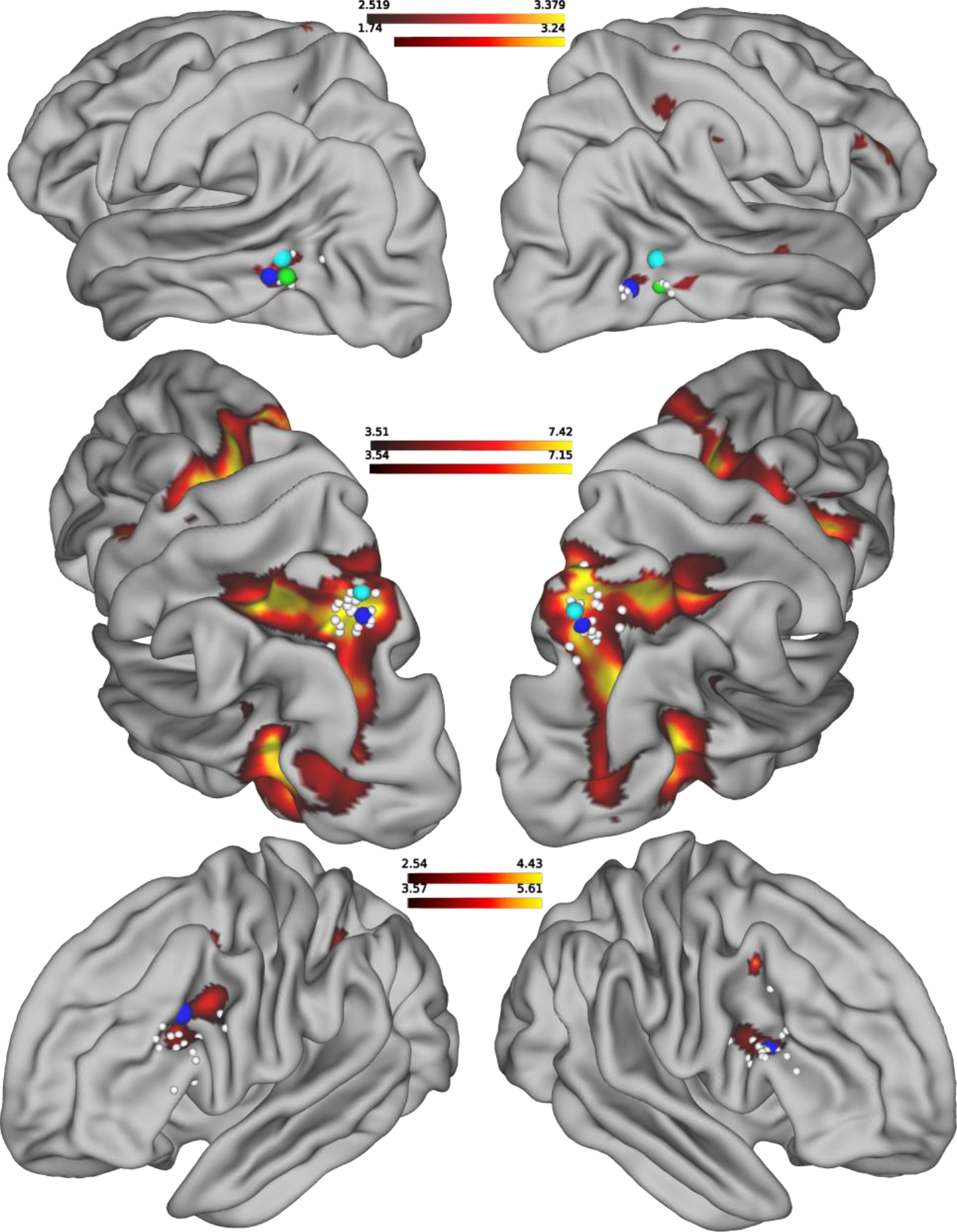
Activation maps. Blue dots mark maximum activation on the group level closest to locations taken from literature (green^11^ and cyan^12^ dots), white dots mark the maximum activation of those locations which were identifiable on the individual level. Upper row: contrast gf > cm (localizer paradigm) used to identify the GFP; Middle row: contrast cm > bl (localizer paradigm) used to identify saccade-related activity in the hLIP closest to location taken from^13^ (cyan dot); Bottom row: uninf > ua (contextual gaze following paradigm).

An analogous procedure was applied to localize the hLIP using the contrast *cm* > *bl*, again based on trials from the first experiment. The location of maximum activation at the group level was found to be at *x*, *y*, *z* = 21, −67, 50 (right) and *x*, *y*, *z* = −21, −67, 53 (left) (blue spheres ibid.) in good accordance with previous work on saccade related activity in the parietal cortex ^13^ (Fig. 3, middle). The generally much stronger contrast allowed us to determine individual contrast hotspots for all participants when considering the aforementioned secondary criteria described except for the proximity criterion (white spheres ibid., SD of individual locations: right x, y, z = 4, 5, 5; left x, y, z = 4, 3, 5). The latter was not considered because of the wide expanse of significant contrast in parietal cortex.

In order to identify brain regions specifically activated when the other’s gaze is not sufficient to unambiguously single out a target object we ran an exploratory whole-brain analysis. Using the BOLD data from the contextual gaze following experiment, we calculated the BOLD contrast between trials from both ambiguous conditions vs. the unambiguous condition. This contrast was significant (*p* <= 0.001, cluster size >= 6 voxel) for a region in the inferior prefrontal cortex (Fig. 3, bottom) whose group level maxima were found in slightly different locations in the two hemispheres, namely at *x*, *y*, *z* = −39, 11, 29 in the left and *x*, *y*, *z* = 48, 20, 23 in the right hemisphere (blue spheres), corresponding to the most lateral part of left BA 8 and the upper right BA 44. In 15 subjects we could delineate individual contrast locations that complied with the criterion of a significant activation of at least six adjacent voxel at a threshold of *p* = 0.05 (white spheres ibid., SD of individual locations: right x, y, z = 5, 6, 6; left x, y, z = 5, 8, 6). The individual locations scattered around BA 44, BA 8 and BA 9 and henceforth we will refer to this region as the inferior frontal junction (IFJ). In the absence of *a priori* expectations based on previous studies we did not exclude individual locations that did not match the proximity criterion.

Weaker, albeit still significant *inf/uninf* > *ua* contrasts were also found in the medial part of left BA 8 at *x*, *y*, *z* = −3, 11, 50, bilaterally in BA 6 at *x*, *y*, *z* = −21, −4, 50 and *x*, *y*, *z* = 24, −1, 50 and at *x*, *y*, *z* = 36, 8, 47 (right hemisphere) not far from the IFJ (cf. Supplementary material Tab. 1). Reversing the contrast, i.e. *ua* > *inf/uninf*, we observed bihemispheric significance within BA 13 (insula), BA 40, within the cingulate cortex (BA 24 and 31) and within BA 7 (all *p* = 0.001, and a minimum of 6 adjacent voxel, cf. Supplementary material Tab. 1). All regions mentioned in the preceding paragraph, even though lighting up in the contrast at the given significance level, did not significantly differentiate between conditions in the following examination of the time courses of the BOLD signals.

### Time course of BOLD signals

Successful gaze following in the contextual gaze following task requires the preceding resolution of the object choice ambiguity. The fact that the IFJ exhibited a significant influence of ambiguity suggests that it might play a role in resolving it. In this case, the influence should be apparent well before the onset of gaze following. In order to test this prediction, we examined the temporal development of BOLD responses associated with the three conditions (*unambiguous*, *ambiguous-informative*, *ambiguous-uninformative*) in the IFJ and the other major task-related areas, the GFP and the hLIP. To this end we determined the individual time courses of the BOLD signal within sphere-shaped ROIs. Whenever the localizer experiment had pinpointed significant individual contrast hot spots, spheres with a radius of 5 mm were centered at the hot spot coordinates. If this was not the case, instead spheres with a radius of 10 mm, centered at the group level location of the respective contrast were deployed. Fig. 4 depicts the baseline corrected time courses of the BOLD signals averaged across participants, separately for the three conditions and the six ROIs. For all ROIs we found a clear modulation of the BOLD signal by the sequence of trial events with significant activity also in later phases of a trial, independent of condition, with one qualification: the signal evoked in *unambiguous* trials in the IFJ was weak at best and confined to a short period following the presentation of the cue. On the other hand, in the other two conditions the signal elicited by the cue was not only much stronger but also much more sustained. As anticipated by the activation maps resulting from experiment 1, the hLIP region showed the overall strongest BOLD signals while those in the GFP and the IFJ were on a lower level. The time course of the BOLD signal in the GFP and the hLIP showed structural similarities. An initial drop after 5 s was followed by two peaks, one after 10 s and another after 15 s (IPS)/16.5 s (GFP). We assume that the first peak is related to the onset of the cue and the second to the go-signal. The BOLD signal in the IFJ exhibited a qualitatively different shape: the signal appeared to rise in response to the cue (clearly only for the two ambiguous conditions) but there was no second peak in relation to the go-signal. To test for significant differences between conditions we performed a permutation test at each time point (FDR corrected). This test yielded significant differences between the *unambiguous* and the *ambiguous-uninformative* condition between 14 s and 17 s in both hemispheres (FDR(*p*) < 0.05) and in the IFJ between 10.6 s and 17 s (left) and 10.6 s and 15.4 s (right) (FDR(*p*) < 0.05) (gray shaded areas in Fig. 4). In other words, the IFJ differentiates earlier between ambiguity condition than the IPS. The profiles for *ambiguous-informative* and the *ambiguous-uninformative* were very close and statistically not different from each other in both the IFJ and the hLIP region.

**Fig. 4.**
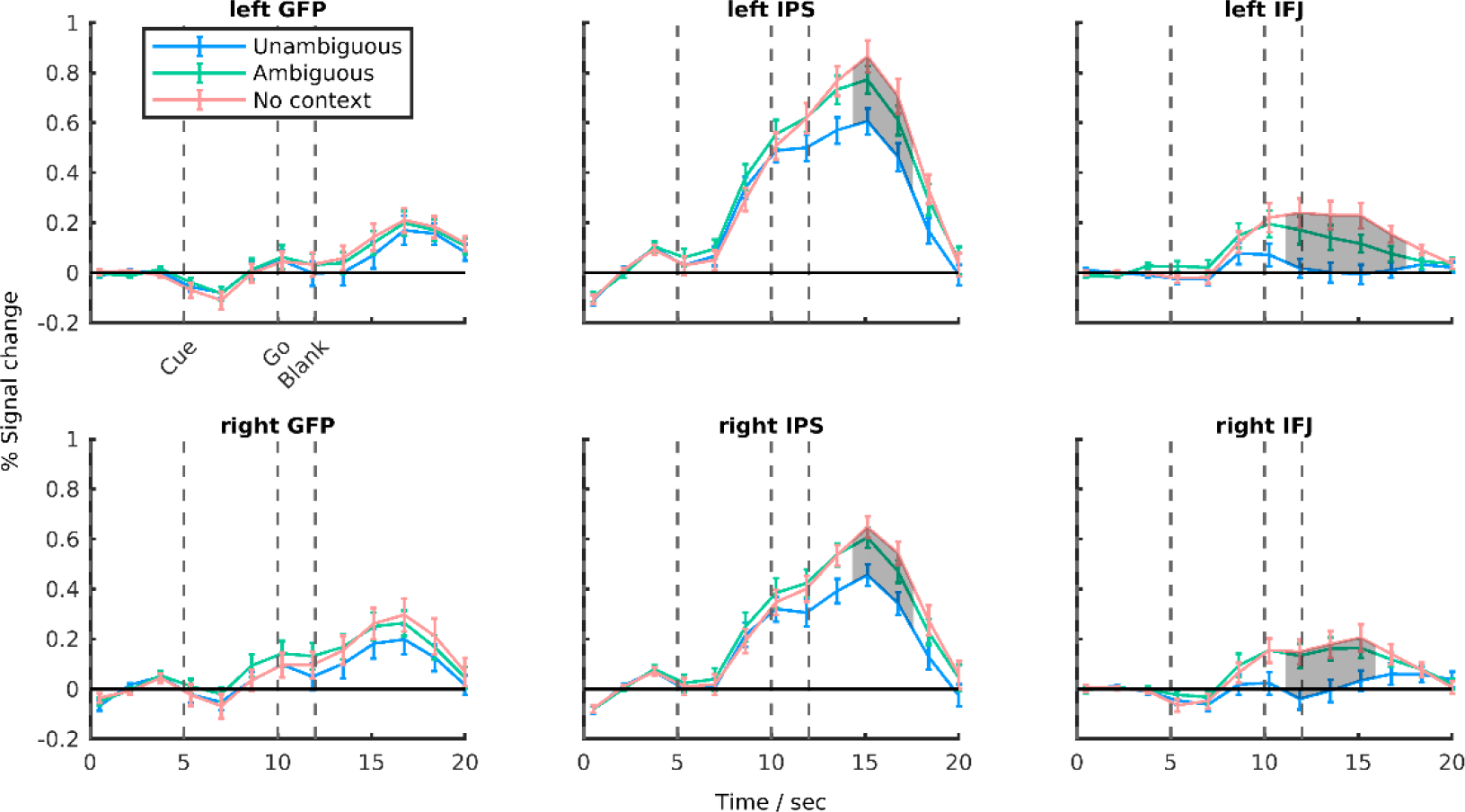
Time courses of activation. Time course of mean percent signal change (error bars are SEM). Areas in which conditions showed significant differences are shaded (permutations test, FDR(p) < 0.05).

Also, the other areas mentioned in the preceding section on task-related brain areas exhibited BOLD signals that showed a modulation by the sequence of task events. Yet, these profiles did not distinguish between conditions.

## Discussion

This study confirms our previous finding that the GFP in the pSTS plays a major role in processing information on the others’ gaze in order to establish joint attention. The present work shows that this role is confined to extracting information on gaze direction. No matter if one or more potential target objects are hit by the gaze vector, the BOLD activity in the GFP is the same. The need to differentiate between objects in case more than one is lying on the gaze vector recruits additional areas that exhibit differential activity. One of these areas, the hLIP in the parietal lobe is also activated in the more traditional, restricted gaze following paradigms, in which the gaze hits one object only. hLIP is necessary for the control of spatial attention^14^. Work on monkey area LIP, arguably homologous to hLIP, has suggested that this area constitutes a priority or saliency map that attracts the “spotlight” of attention to a highlighted map location. The highlighting may be a consequence of bottom-up sensory cues, of symbolic cues or of gaze cues^15,16^. The latter is suggested by single unit recordings from area LIP. Many LIP neurons respond to the appearance of a gaze cue provided the gazed at location lies within the neuron’s receptive field^17^. Spatial selectivity for gazed at locations and objects at these locations is also exhibited by many neurons in monkey GFP^18^. However, unlike neurons in LIP, those in the GFP are selective for gaze direction cueing and do not respond to bottom-up sensory cues highlighting a specific spatial location. This selectivity suggests that the priority map in LIP might draw on input from the GFP. The yoked activation of the hLIP/LIP and the GFP in BOLD imaging studies of gaze following is in principle in accordance with this scenario^11,12,17^. However, the poor temporal resolution of the BOLD signals does not allow us to critically test if the assumed direction of information flow holds true. In any case, bidirectional projections are known to connect monkey area LIP and parts of the STS^19^. One well-established pathway links area LIP and PITd, an area in the lower STS, probably close to the GFP, known to contribute to the maintenance of sustained attention^20,21^. Yet, the anatomical data available does not allow us to decide if the GFP may indeed be contributing to this fiber bundle.

The BOLD signal evoked by gaze following in the hLIP was overall much stronger than in the GFP. Moreover, unlike the GFP signal, it exhibited a clear dependence on the condition. Higher activity was associated with the *ambiguous-informative* and the *ambiguous-uninformative* conditions, both associated with unresolved uncertainty as to the correct object. Why should a region thought to coordinate spatial shifts of attention show an influence of target ambiguity, i.e. the need to choose between several potential targets? One possible answer may be that the higher hLIP activity reflects an increased attentional load. More specifically, increased uncertainty in ambiguous trials may have prompted more covert shifts of attention from one object to the other in an attempt to resolve the ambiguity. Although we found no difference in the number of exploratory saccades after the go signal across conditions, we cannot rule out that participants covertly shifted attention between targets in ambiguous trials more than in the other trials and that this might have led to the observed increased activity in the area hLIP. However, a more parsimonious explanation could be that the hLIP constitutes a neural substrate for making decisions under uncertainty independent of the attentional load as suggested by several studies such as^22^.

A qualitatively similar dependency on condition also characterized BOLD activity in a region we identified as IFJ based on its location in the frontal lobe at the junction between premotor cortex (BA 6), BA 44 and BA 8. The condition dependency of the IFJ signal is most probably a consequence of the need to shift attention between the two object categories, houses and hands. This interpretation draws on an MEG-fMRI study carried out by Baldauf and Desimone that demanded the allocation of attention to distinct classes of visual objects such as faces and spatial scenes^23^. Depending on the object of attention, gamma band activity in the IFJ was synchronized either with the fusiform face area (FFA) or the parahippocampal place area (PPA).

Hence, the IFJ seems to play a role in allocating attention between objects or object categories and shifting between items. Related work on the putative monkey homologue of human IFJ, the ventral pre-arcuate (VPA), suggests that object representations become highlighted by a match of object templates in VPA and vision-based object representations in inferotemporal cortex^24^. Arguably, the need to choose an object in the ambiguous conditions in our experiment requires a deeper scrutiny of the object options in order to find the match with the object template. This increased effort may be the cause of the stronger IFJ BOLD signal associated with the ambiguous conditions. Within this framework, IFJ can be assumed to highlight specific object representations in inferotemporal cortex. If this was true, information needed by the hLIP to disambiguate the object choice for gaze following would have to be tapped from inferotemporal cortex rather from the IFJ.

In sum, our results suggest a fronto-temporo-parietal network for gaze following and the allocation of joint attention underlying the disambiguation of object choices if more than one object is met by the other’s gaze vector. Information on the direction of the other’s gaze is provided by the GFP, information that allows the hLIP to highlight the spatial positions of all objects lying on the gaze vector. Object-based attention, guided by the IFJ, highlights a relevant object category. The intersection between the two will substantially reduce the possible choices, in most cases singling out just one object that then will become the target of the observer’s gaze following response, elicited by the hLIP.

## Methods

### Participants

Nineteen healthy, right-handed volunteers (9 females and 10 males, mean age 27.4, s.d. = 3.6) participated in the study over three sessions. Participants gave written consent to the procedures of the experiment. The study was approved by the Ethics Review Board of the Tübingen Medical School and was carried out in accordance with the principles of human research ethics of the Declaration of Helsinki.

### Task and procedure

The study was conducted in three sessions across separate days. On day 1, we instructed participants about the study goals and familiarized them with the experimental paradigms outside the MRI-scanner by carrying out all relevant parts of the fMRI experiments. The following fMRI-experiments included a functional localizer paradigm for the scanning session on day 2 as well as a contextual gaze following paradigm for the scanning session on day 3.

#### Behavioral session

After participants had been familiarized with the tasks, they were head-fixed using a chinrest and a strap to fix the forehead to the rest. Subjects were facing towards a frontoparallel screen (resolution = 1280 × 1024 pixels, 60 Hz) (distance to eyes ≈ 600 mm). Eye tracking data were recorded while participants had to complete 80 trials of the localizer paradigm and 72 trials of contextual gaze following.

#### Localizer task

We resorted to the same paradigm used in^11^, to localize the gaze following network and in particular its core, the GFP. In this paradigm, subjects were asked to make saccades to distinct spatial targets based on information provided by a human portrait presented to the observer. Depending on the instruction, subjects either had to rely on the seen gaze direction to identify the correct target (*gaze following* condition) or, alternatively, they had to use the color of the irises, changing from trial to trial but always mapping to one of the targets, in order to make a saccade to the target having the same color (*color mapping* condition). In other words, the only difference between the two tasks was the information, subjects had to exploit in order to solve the task, while the visual stimuli where the same.

This task is associated with higher BOLD activity in the GFP, a region, close to the pSTS, when people perform gaze following compared to color mapping. The task is further associated with the activation of regions in the intraparietal sulcus (IPS) as well as the frontal cortex that take part in controlling spatial attention and saccade generation^11,12^. Out of the 19 subjects of our study, 16 performed 6 runs (40 trials per run) and for reasons of time management during image acquisition, one subject performed 5 runs and two subjects performed 4 runs.

#### Contextual gaze following task

An example of a trial is shown in Fig. 1. Each trial consisted of the following events in sequence. The trial started by or with the appearance of an avatar (size in angular deg.) image in the center of the screen together with four arrays of drawn objects (houses and hands, 3 objects per array). Subjects were asked to fixate on a red fixation dot (diameter) between the portrait’s eyes. After 5 seconds of baseline fixation, the portrait’s gaze shifted towards one specific target object. Simultaneously, an auditory contextual instruction either specified the object class of the target (spoken words “hand” or “house”) or was not informative (“none”). While maintaining fixation, subjects needed to judge which object the target was (i.e. on which object the face was most likely looking at). After 5 seconds delay, the fixation dot vanished, an event that served as a go signal. Participants had 2 seconds to make a saccade to the chosen target object and fixate it until a subsequent blank fixation screen was presented for 8 seconds. The subjects were instructed to perform the task as accurate as possible. They were specifically instructed, when unsure about the actual target, they should still rely on gaze and contextual information and choose the target they believed the avatar to be looking at.

### Stimuli

Control of visual and auditory stimuli as well as data collection was controlled by the Linux based open source system *nrec (https://nrec.neurologie.uni-Tübingen.de/)*. The stimuli in the localizer task were identical to the stimuli used in a previous study^11^. The stimuli of the contextual gaze following task consisted of an avatar and in total 12 target objects belonging to different types (houses and hands). The avatar was generated with the custom-made OpenGL library *Virtual Gaze Studio*^25,26^ which offers a controlled virtual 3D-environment in which an avatar can be set to precisely gaze at specific objects. More specifically, the program allows to place objects on a circle, parallel to the coronal axis, anterior to the avatar face. For each stimulus, we placed 12 objects in the surroundings of the avatar. The location of individual objects was fully determined by the distance to the coronal plane at the level of the avatar’s nasion, the radius of the circle and the angle of the object on that circle. By keeping the angle on the circle constant for sets of three objects, we created four arrays at angles 120°, 150°, 210° and 240°. The individual locations of these objects were specified by varying the distance and the circle radii based on trigonometric calculations. For these calculations we assumed a right triangle from the avatar’s nasion with the hypotenuse pointing towards the object, an adjacent leg (length corresponded to the distance of the circle) proceeding orthogonal to the coronal plane, and an opposite leg which corresponded to the radius. By keeping tan𝛼 fixed to 0.268, we varied the distances and circle radii. For the 120° and 240° arrays, the circle radii were 335, 480, 580 and the distances were 90, 129 and 151 virtual mm. For the 150° and 210° arrays, the radii were 380, 510 and 590 and the distances were 102, 137 and 158 virtual mm. The reason for the difference of radii and distances between 120°/240° and 150°/210° arrays was that this allowed to exploit the total width of the screen. This procedure guaranteed that the angle of the gaze vector to all objects on an array was almost identical. This makes it relevant to take contextual information into account in order to choose the true target.

The objects were drawings of the two categories houses and hands, downloaded from freely available online sources (http://www.allvectors.com/house-vector/, https://www.freepik.com/free-vector/hand-drawn-hands_812824.htm#term=hands&page=1&%20position=37). The target objects were arranged in four radial directions (three objects in each direction) with the avatar eyes as the origin; in other words, the avatar’s gaze always hit one out of three objects along the gaze vector though participants were not able to tell which of the three it was. On each array, either 2 hands and one house or one hand and two houses were present. Further, we fixed the number of hands and houses per hemifield to three. The relative order of the objects was pseudo-randomized from trial to trial.

During a trial the participant observed the avatar making a saccade in one of the four directions while simultaneously hearing a verbal instruction providing the additional information by either specifying the target type (“house” or “hand”) or being uninformative in that respect (“none”) (cf. Fig. 1 for an illustration). In connection with the set of targets specified by the gaze cue the verbal instruction created different levels of ambiguity: *unambiguous* (only one of the verbally specified types was in the set), *ambiguous-informative* (two of the types were in the set) and *ambiguous-uninformative* (verbal instruction was uninformative, i.e. three possible targets). We created a pool stimulus sets which satisfied three constraints: There was an equal number of trials in which a) the targets were hands or houses, b) targets were presented with an *unambiguous*, *ambiguous-informative* and *ambiguous-uninformative* instruction, and c) the spatial position (one out of twelve potential positions) of targets was matched. This led to 2 × 3 × 12 = 72 stimuli sets. We exposed every subject to 180 trials in which each stimulus set was shown twice and for the residual 36 trials, stimuli were drawn from pseudo-randomly from the stimulus pool so that the three criteria above were met.

Auditory stimulation was delivered via headphones (Sennheiser HD 201, Wedemark-Wennebostel, Germany, during the behavioral session and the standard air pressure headphones of the scanner system during the MRI sessions). The auditory instructions “hand”, “house” and “none” were computer generated with the web application imTranslator (http://imtranslator.net/translate-and-speak/speak/english/) and processed with the software Audacity 2.1.2. The sound files had a duration of 600 ms.

### Eye tracking

During all three sessions, we recorded eye movements of the right eyes using commercial eye tracking systems (Behavioral sessions: Chronos Vision C-ETD, Berlin, Germany, sampling rate 400 Hz, resolution < 1° visual angle; Scanning sessions: SMI iView X MRI-LR, Berlin, Germany, sampling rate = 50 Hz, resolution ≈ 1° visual angle).

Eye tracking data was processed as follows. First, we normalized the raw eye tracking signal by dividing it by the average of the time series. Eye blinks were removed using a velocity threshold (> 1000 °/s visual angle). Next, we focused on a time window in which we expected the saccades to the target objects to occur ([go-signal – 500 ms, go-signal + 1800 ms]). Within this time window, we detected saccades by identifying the time point of maximum eye movement velocity. Pre- and post-saccadic fixation positions were determined by averaging periods of 200 ms before and after the saccade occurred. Due to partly extensive measurement noise of the eye tracking system, we did not automatize the categorization of the final gaze position. Instead, we plotted X- and Y coordinates of the post-saccadic eye position for every run. An investigator (MG), who was blind to the true gaze target-directions of the stimulus face, manually validated, which trials yielded positions that were clearly assignable to one object location. For the behavioral analysis we only used the valid trials (mean number of valid trials per participant = 80.2, s.d. = 45.4, range = [0, 153]) and weighted the individual performance values by its number in order to compute weighted means and SDs. Note, that we used these valid trials only for the behavioral analysis but used all trials of the participants for the fMRI analysis, assuming that eye tracking measurement noise was independent of the performance of the subjects.

### fMRI acquisition and preprocessing

We acquired MR images using a 3T scanner (Siemens Magnetom Prisma, Erlangen, Germany) with a 20-channel phased array head coil at the Department of Biomedical Magnetic Resonance of the University of Tübingen. The head of the subjects was fixed to the head coil by using plastic foam cushions to avoid head movements. An AutoAlign sequence was used to standardize the alignment of images across sessions and subjects. A high-resolution T1-weighted anatomical scan (MP-RAGE, 176 × 256 × 256 voxel, voxel size 1 × 1 × 1 mm) and local field maps were acquired. Functional scans were carried out using a T2^*^-weighted echo-planar multi-banded 2D sequence (multi-band factor = 2, TE = 35 ms, TR = 1500 ms, flip angle = 70°) which covered the whole brain (44 × 64 × 64 voxel, voxel size 3 × 3 × 3 mm, interleaved slice acquisition, no gap). For image preprocessing we used the MATLAB SPM12 toolbox (Statistical Parametric Mapping, https://www.fil.ion.ucl.ac.uk/spm/). The anatomical images were segmented and realigned to the SPM T1 template in MNI space. The functional images were realigned to the first image of each respective run, slice-time corrected, coregistered to the anatomical image. Structural and functional images were spatially normalized to MNI space. Finally, functional images were spatially smoothed with a Gaussian kernel (6 mm full-width at half maximum).

### fMRI analysis

We estimated a generalized linear model (GLM) to identify ROIs of single subjects. On these regions, we performed time course analyses to investigate event-related BOLD signal changes. In a first-level analysis, we constructed GLMs for the localizer task (GLM_loc_) and the contextual gaze following task (GLM_cgf_). The GLM_loc_ included predictors at the onsets of directional cues and of the baseline fixation phase. The GLM_cgf_ had predictors at the onset of the contextual instruction. These event specific predictors of both GLMs used the canonical hemodynamic response function of SPM to model the data. We corrected for head motion artifacts by the estimation of six movement parameters with the data of the realignment preprocessing step. Low-frequency drifts were filtered using a high-pass filter (cutoff at 1/128 Hz).

### GFP and hLIP localizer

Before collecting the data, we specified the expected locations of two brain areas, hLIP and GFP from fMRI literature. We resorted to the hLIP coordinates of the human homologue of monkey area LIP which had been identified in humans who performed a delayed saccade task^13^. We transformed the coordinates into MNI space, using an online transformation method of Lacadie and colleagues^27^ (http://sprout022.sprout.yale.edu/mni2tal/mni2tal.html). ROIs were defined as the voxel of highest signal contrast (GLM_loc_: directional cue vs. baseline fixation) the cluster of significant activity (cluster size ≥ 6, p < 0.05) which minimized the spatial distance to the standard coordinates. This contrast has been associated with shifts of attention in response to gaze cues (Marquardt, Ramezanpour et al. 2017). We identified the hLIP regions bilaterally in all 19 subjects with a mean distance of 13.4 mm (s.d. = 3.9 mm) between IPS_right_ and the standard coordinates and 11.93 mm (s.d. = 3.7 mm) for IPS_left_. At the location of the ROI, a sphere (radius = 5 mm) was placed.

We used a similar procedure for the GFP but with different expected coordinates, a different contrast of the (GLM_loc_ gaze following vs. color mapping) and the additional constraint that the cluster of significant activity had to be at least partially located within 10 mm distance around the pSTS standard coordinates. This contrast has been associated to the calculation of the gaze vector direction (for more details see Marquardt et al., 2017). We localized pSTS_right_ in nine individual subjects (mean distance = 6.6 mm, s.d. = 3.1 mm) and pSTS_left_ in six subjects (mean distance = 7.7 mm; s.d. = 1.4 mm). For those subjects and hemispheres where we did not identify pSTS, we reasoned that signal contrast was not high enough and therefore placed a sphere (radius 10 mm) at the coordinates obtained from a second level analysis.

### Contextual gaze following analysis

We performed an exploratory whole-brain analysis on the data from the contextual gaze following task. We contrasted ambiguous conditions with the unambiguous condition at the group level (significance threshold p < 0.001, cluster size >=6 voxel) as well as at the single subject level (significance threshold p < .05, cluster size ≥ 6 voxel). For the single subject analysis, we searched for ROIs that minimized the distance to the group level coordinates. At the identified individual locations (15 subjects) we placed spheres of 5 mm radius. Again, we used 10 mm spheres at the group level coordinates for those four subjects for whom we had not identified the ROI in the first level analysis.

For every ROI, the mean raw time series of the BOLD signal was extracted using the MATLAB toolbox *marsbar 0.44* (http://marsbar.sourceforge.net). The time course of every trial was normalized by the average signal intensity 5 s before the contextual instruction onset and transformed into % of signal change. For each participant, we averaged time courses across trials and used the time courses of the three contextual conditions and six ROIs for our analysis. To test differences across conditions for statistical significance, we performed permutation tests at each time point after contextual instruction delivery. To do so we pooled the data of two experimental conditions, respectively, and produced 10,000 random splits for each pool. By computing the differences between the means of these splits, we obtained a distribution of differences under the null hypothesis. Calculating the fraction of values more extreme than the actual difference between means allowed us to obtain a *p*-value for each time bin. To account for the multiple comparison problem, we transformed *p*-values to FDR corrected *q*-values^28^ and considered each time bin with *q* < .05 as statistically significant.

**Table 1.**
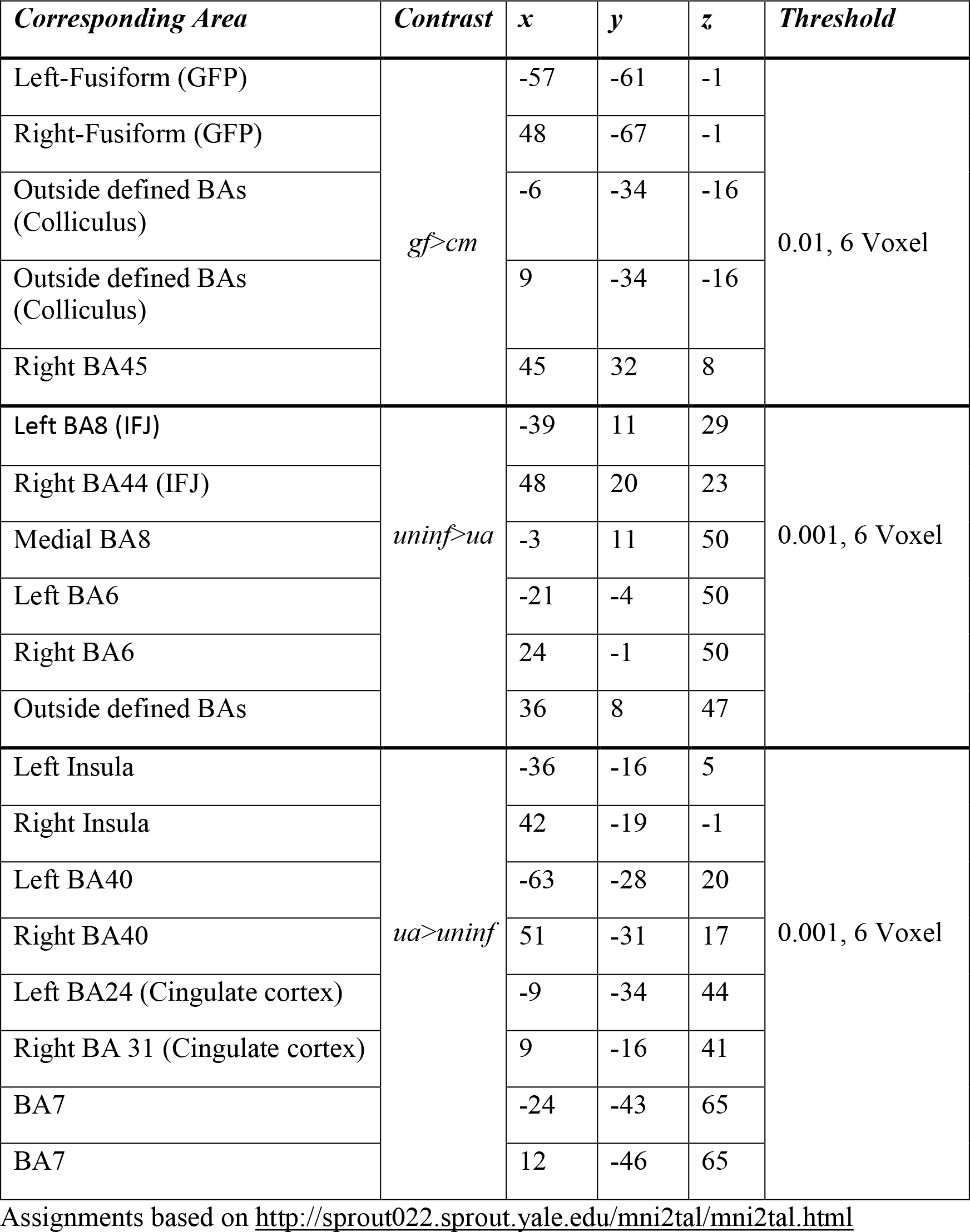

## Acknowledgments

We are grateful to Friedemann Bunjes and Michael Erb for technical support. This work was made possible by a grant from the Deutsche Forschungsgemeinschaft (TH 425/12-2).

## Authors Contributions

PT developed the conceptual framework of the research. PT, PK and HR designed the experiments. PK and PWD performed the experiments. PK and MG analyzed the data. All authors contributed to the interpretation of results and the writing.

## Competing Interests statement

All authors declare to have no competing interests of any sort.

## Supplement

### Localizer experiment

As a localizer task we used a cued saccade task, also denoted as a *gaze following vs. color mapping* task^11^. During a baseline fixation phase, subjects had to fixate on a red dot between the eyes of a photography of a face gazing straight ahead. Below the stimulus face, five colored and horizontally arranged rectangles were presented as gaze targets. After five seconds of baseline fixation, the portrait’s eye-gaze shifted towards one of the targets and, simultaneously, its eye color (i.e. the color of the irises) changed to match the color of one of the rectangles. After one second, the red dot disappeared (go signal) and the subjects had to shift their own gaze towards to the correct target and fixate it. There were two different experimental conditions: (1) in *gaze following* trials, the correct target was determined by the eye-gaze direction of the stimulus face, (2) in *color mapping* trials, the correct target had the same color as the stimulus irises. The task was performed in several runs, each consisting of four blocks (2 gaze following, 2 color mapping). Each block started with the task instruction as a seven seconds lasting window containing the written words “gaze following” or “color mapping”, followed by 10 corresponding trials. Task instruction alternated between blocks. Target objects were counter-balanced such that each rectangle was the target object twice during a block and target order was pseudorandomized.

